# Apoptotic cells induce CD103 expression and immunoregulatory function in myeloid dendritic cell precursors through integrin α_v_ and TGF-β activation

**DOI:** 10.1101/2020.04.14.040923

**Authors:** Ailiang Zhang, Helena Paidassi, Adam Lacy-Hulbert, John Savill

**Affiliations:** Medical Research Council Centre for Inflammation Research, Queen’s Medical Research Institute, University of Edinburgh, Edinburgh BioQuarter, Edinburgh, UK; Inserm, U1111, Lyon, France; Benaroya Research Institute at Virginia Mason, Seattle, USA

## Abstract

In the mammalian gut CD103+ve myeloid DCs are known to suppress inflammation threatened by luminal bacteria, but stimuli driving DC precursor differentiation towards this beneficial phenotype are incompletely understood. We isolated CD11+ve DCs from mesenteric lymph nodes (MLNs) of healthy mice; CD103+ve DCs were 8-24 fold more likely than CD103-ve DCs to exhibit extensive of prior phagocytosis of apoptotic intestinal epithelial cells. However, CD103+ve and CD103-ve MLN DCs exhibited similar *ex vivo* capacity to ingest apoptotic cells, indicating that apoptotic cells might drive immature DC differentiation towards the CD103+ve phenotype. When cultured with apoptotic cells, myeloid DC precursors isolated from murine bone marrow and characterised as lineage-ve CD103-ve, displayed enhanced expression of CD103 and β8 integrin and acquired increased capacity to induce Tregs after 7d *in vitro.* However, DC precursors isolated from α_v_-tie2 mice lacking α_v_ integrins in the myeloid line exhibited reduced binding of apoptotic cells and complete deficiency in the capacity of apoptotic cells and/or latent TGF-β1 to enhance CD103 expression in culture, whereas active TGF-β1 increased DC precursor CD103 expression irrespective of α_v_ expression. Fluorescence microscopy revealed clustering of α_v_ integrin chains and latent TGF-β1 at points of contact between DC precursors and apoptotic cells. We conclude that myeloid DC precursors can deploy α_v_ integrin to orchestrate binding of apoptotic cells, activation of latent TGF-β1 and acquisition of the immunoregulatory CD103+ve β8+ve DC phenotype. This implies that a hitherto unrecognised consequence of apoptotic cell interaction with myeloid phagocytes is programming that prevents inflammation.

## INTRODUCTION

Mammalian bowel contents are laden with potently pro-inflammatory bacteria and yet the healthy gut is not inflamed. There is now evidence that gut inflammation can be kept in check by anti-inflammatory programming of myeloid phagocytes encountering cells dying by apoptosis in the gut wall. Elegant work from Blander’s group employed transgenic techniques to induce apoptosis in intestinal epithelial cells (IECs) *in vivo* and then track their uptake by myeloid phagocytes. Dendritic cells (DCs) expressing CD103 were observed to ingest apoptotic IECs and respond with two notable patterns of gene transcription; downregulation of inflammatory genes and suppression of the immune response [1]. Previous studies had reported that CD103+ve myeloid DCs in the gut wall and draining lymphatics directed powerful suppression of immune responses, inducing peripheral T regulatory lymphocytes (Tregs) in draining mesenteric lymph nodes through mechanisms including TGF-β1 activation [2–6]. However, this body of work had not addressed whether CD103+ve myeloid DCs in the gut had interacted with cells spontaneously undergoing apoptosis in the gut wall. Nevertheless, murine CD103+ve myeloid DCs in the mouse gut have been considered analogous to OX41-ve DCs in rats, which are known to ingest IECs spontaneously undergoing apoptosis and transport these to T cell areas of mesenteric lymph nodes in a manner suggesting a role in the induction and maintenance of self-tolerance [7].

There is growing evidence that myeloid DC α_v_ integrins are crucial for capture of apoptotic cells and associated anti-inflammatory TGF-β1 activation. Thus, α_v_ β_5_ mediates myeloid DC recognition of apoptotic cells [8–10] and α_v_ β_8_ is crucial for binding and activation of latent TGF-β1[3, 11–13]. When we deleted α_v_ in the myeloid line, knockout mice housed in conventional SPF conditions, but known to be colonised by *Helicobacter spp.*, spontaneously developed ulcerative colitis associated with a reduction in Tregs in the colon and diminished *ex vivo* capacity of gut-associated myeloid DCs to ingest apoptotic cells efficiently and induce Tregs. Furthermore, there was a reduction in CD103+ve CD11c+ve DCs in mesenteric lymph nodes of mice lacking α_v_ in the myeloid line, suggesting that α_v_ might be necessary for the development of CD103+ve DCs from putatively CD103-ve precursors, although this possibility was not dissected further [9].

In this study we sought evidence that apoptotic cells arising spontaneously in the healthy mouse gut are instrumental in inducing the CD103+ immunoregulatory phenotype in DCs migrating to mesenteric lymph nodes (MLNs). In keeping with this hypothesis, we report that in healthy mice, CD103+ve MLN DCs were around 8-fold more likely to exhibit histochemical evidence of prior spontaneous ingestion of apoptotic IECs than CD103-ve MLN DCs, despite the two DC subpopulations having similar capacity for phagocytosis of apoptotic targets *ex vivo.* In a model system *in vitro*, co-culture with apoptotic cells promoted differentiation of lineage-ve CD103-ve myeloid DC precursors towards an immunoregulatory phenotype, with increased expression of CD103 and the β_8_ integrin chain necessary for TGF-β1 activation, coupled with increased capacity to induce Tregs. Immunostaining demonstrated that α_v_ integrin and latent TGF-β1 co-localised at points of contact between apoptotic cells and DC precursors. Furthermore, by contrast with wild type, DC precursors prepared from mice defected for α_v_ in the myeloid line failed to increase CD103 expression when co-cultured with apoptotic cells and/or latent TGF-β1, but did do so when cultured with active TGF-β1.

We conclude that immature myeloid dendritic cell precursors can deploy α_v_ integrins to capture apoptotic cells and co-ordinately activate latent TGF-β1 with the result that such precursors differentiate towards the CD103+ve immunoregulatory DC phenotype. Such DCs are known to be crucial in suppression of inflammation in the murine gut [2–6] and the current study confirms that CD103+ve DCs migrating to mesenteric lymph nodes exhibit extensive evidence of prior ingestion of apoptotic cells arising spontaneously in the gut wall. These data suggest that future studies should examine whether tonic suppression of inflammation in the gastrointestinal tract is a newly identified function for clearance of apoptotic cells in addition to those previously described in resolution and repair of established inflammation [14–16].

## Materials and Methods

### Animals

Female C57BL/6 mice (6-8 weeks old) were purchased from Charles River laboratories UK. Foxp3+eGFP mice and CD45.1 OT-II mice were generously provided by Stephen Anderton’s Group (MRC Centre for Inflammation Research, University of Edinburgh). We have previously demonstrated [9] targeting of α_v_ deletion in α_v_-tie2 mice backcrossed to C57BL/6 backgrounds for 10 generations. In these experiments control mice were wild type genotype from the same litters at 6 to 8 weeks old. All mice were housed in conventional specific and opportunistic pathogen-free facilities. All experiments were performed in accordance with the UK Home Office Scientific Procedures Act (1986) and local ethical approval.

### Isolation of CD103+ve DC and CD103-ve DC from MLN and immunofluorescence microscopy

Mesenteric lymph nodes (MLN) were collected from euthanised naïve female C57BL/6 mice and were digested in RPMI1640 containing DNaseI (20 mg/ml) and Liberase TL (0.33 mg/ml; both from Roche) for 30 minutes and then incubated in PBS/ 2% BSA/ 10 mM EDTA for another 5 minutes. Cells were filtered in PBS/ 2% BSA/ 2 mM EDTA by passing through 40µm cell strainers and CD11+ve cells were enriched by CD11c-MACs beads (Miltenyi Biotec). CD11c+ CD103+ve and CD103-ve dendritic cells were further sorted by flow cytometry [3]. The sorted CD103+ve and CD103-ve DCs cells were used for fluorescent staining.

For apoptotic cell DNA detection in MLN DC, the FACS-sorted CD103+ve and CD103-ve DC were processed by the TUNEL (terminal deoxynucleotidyl transferase–mediated dUTP-biotin nick-end labelling) method using an TACS™TdT In-Situ Apoptosis Detection Kit (R&D) according to the manufacturer’s instructions and nuclei subsequently labelled with DAPI.

For detection of NSE, the FACS-sorted single cell suspensions were prepared by cytospin and fixed with 2% paraformaldehyde. NSE reactivity was developed by using α-naphthyl acetate (Sigma) as substrate. All images were obtained using an Axioskop microscope (Zeiss Ltd, UK).

### Generation of CD103+ve and CD103-ve DCs from myeloid DC precursors

DC precursors were isolated from C57BL/6 mouse bone marrow as described previously [17] with minor changes. Briefly mouse bone marrow cells were harvested by flushing femurs and tibias and passing through a 40µm cell strainer. The cells were stained with a biotin conjugated cocktail of lineage-specific Abs (CD3, CD11c, CD45R, NK1.1, Ter-119, ly6G; from BD bioscience) and negatively selected by anti-biotin microbeads (Miltenyi Biotec). These untouched lineage negative cells were CD11c-ve CD103-ve, considered to be DC precursors, and were cultured in 24 well plates at 0.5×10^6^ cells/ml in RPMI 1640 supplemented with 2 mM L-glutamine, 100 U/ml penicillin, 100 µg/ml streptomycin, 10% heat-inactivated FBS, 50µm β-ME and 20 ng/ml GM-CSF (From Invitrogen) or in serum free X-VIVO 15 medium (LONZA) with 20ng/ml GM-CSF. Recombinant human active-TGF-β1 (0.2 ng/ml) and purified human latent TGF-β1 (2 ng/ml) (all R&D Systems) and apoptotic cells were added in at the indicated time points and ratios as described in Figure legends. To block active TGF-β1, cells were cultured in presence of anti-TGF-β antibody at 2μg/ml (1D11; R&D Systems). Cells were fed every two to three days with fresh medium and were harvested at day 6 to 8.

### Generation of apoptotic cells

Human neutrophils were extracted from peripheral blood of healthy volunteers, as described previously [14]. After overnight aging, the cells were routinely 70 – 80% apoptotic, assessed by positive Annexin V and negative propidium iodide (PI) staining; <5% were PI+ve. Mouse thymus was removed and disaggregated by passing through a 40um cell strainer in RPMI 1640 supplemented with 2 mM L-glutamine, 100U/ml penicillin, and 100g/ml streptomycin, to yield a single-cell suspension. Thymocytes were aged overnight at 4×10^6^/ml in RPMI 1640 supplemented medium with 1% FBS and 1µm of Dexamethasone, which yielded a largely annexin V-positive (>85% ± 5%) and low PI-positive (<15% ±5%) cell population [9, 12, 15].

### *In vitro* assay of binding apoptotic cells

DC precursors from bone marrow were labelled with CM-orange (Invitrogen) and cultured with fluorescently labelled (Cell Tracker Green CMFDA, Invitrogen) apoptotic cells at a ratio of 1:5 (DC:AC) for two or three hours as indicated in figure legends. Cells were harvested and analyzed by flow cytometry.

For fluorescence microscopy, cells were fixed with 4% paraformaldehyde and images were obtained using Axioskop microscope (Zeiss Ltd, UK).

### Flow cytometry and antibodies

The single cell suspensions were pre-incubated with 5µg/ml blocking antibody against CD16/CD32 (eBioscience) to reduce nonspecific binding. The cells were stained in PBS containing 0.2% BSA and 0.02% NaN3 with the following antibodies: anti-CD11c-APC (HL3), anti-CD103-PE (M290), B220-PE (all from BD bioscience), MHC II-Brilliant Violet 421 (Biolegend), anti-CD11b-Per-CP-Cy5.5 (M1/70) and mouse regulatory T cell staining Kit (PE Foxp3 FJK-16s, FITC CD4, APC CD25) (all from ebioscience). FACS data acquisition was performed on a LSRII (BD Bioscience) running FACS Diva software. FlowJo software (Tree Star, Ashland, OR) was used for data analysis. Cell sorting of CD103+ve and CD103-ve DC was performed on a FACSAria flow cytometer (BD Bioscience).

### *In vitro* FoxP3+ve T regulatory cell generation

Lymphocytes were harvested from spleens of FoxP3-GFP mice by physical disruption. CD4+ve T cells were pre-sorted by magnetic separation (CD4+ pre-enrichment kit, Stem Cell Technologies). Naïve CD4+ve FoxP3-ve GFP-ve T cells were further sorted by FACs. CD11c+ve BMDCs were harvested from cultures with apoptotic cells or without as control and resuspended into 1×10^6^/ml and cultured with naïve CD4+ve FoxP3-ve T cells in serum free X-VIVO 15 medium (LONZA) supplemented with 2 mM L-glutamine, 100 U/ml penicillin, 100 μg/ml streptomycin, 50 μM 2-β-mercaptoethanol. Cells were co-cultured in the ratio of 1:2.5 (DC: T cell) in the presence of 0.5 μg/ml anti-CD3 antibody (145-2C11; BD Biosciences). Additional cytokines used were recombinant active-TGF-β1 (0.5 ng/ml) and purified human latent TGF-β (2 ng/ml) (R&D Systems) and retinoic acid (100nm, SIGMA). Cells were cultured for 5 days and Treg generation were assessed by GFP expression with flow cytometry.

### *In vitro* Treg functional assay

CD4+ve T cells were isolated from CD45.1 OT-II mouse spleen by negative selection using the CD4+ T-cell isolation kit (Miltenyi Biotec). A biotinylated anti-CD25 antibody was added to the cocktail Ab mixture for depletion of FoxP3+ve CD25+ve T cells. Naïve CD4+ve CD25-ve T cells were labelled with e670 (eFluor® 670, eBioscience) according to the manufacturer’s instructions. These e670 labelled T cells (50×10^3^/well) co-cultured with containing induced regulatory T cells at 1:1 ratio in a round bottom 96 well plates and stimulated by pOVA_323-339_ (from SIGMA) pulsed (1µg/ml for 5hrs) DC (5×10^3^/well). After 72hrs, cells were harvested and stained with CD4-FITC and CD45.1-BV421 (Biolegend). Cell division of e670-labelled CD4+ve T cells was analysed by flow cytometry.

### Quantification of gene expression

BMDC were harvested at day7 and washed with PBS. Then cells were spun down and put into TRIZOL (Invotrogen) for RNA extraction. cDNA was synthesized using High Capacity Reverse Transcription kits (Applied Biosystem). For real-time PCR, all Primer and probes mixes and Taqman fast master mixes were purchased from Applied Biosystems. Gene mRNA expression were quantified by using ABI 7500 fast real-time PCR system according to manufacturer’s instructions. The mRNA relative expression level of target gene was normalized to housekeeping gene 18s.

### Integrin-α_v_ and LAP (latent TGF-β 1) co-localization staining and microscopy

DC precursors from bone marrow cultured with apoptotic cells at a ratio of 5:1 (Ac:DC) for three hours in the presence of GM-CSF (20ng/ml). Cells were harvested and stained with biotin conjugated anti-mouse integrin-α_v_ (BD Bioscience) and goat anti-human LAP (latent TGFb1) (R&D Systems) on ice for 45 minutes. Cells were washed with PBS and stained with secondary antibody streptavidin Alexa 488 and anti-goat Alexa 546 on ice for 20 minutes. Cells were fixed with 4% paraformaldehyde and counter stained with DAPI. The Images were obtained using an Axioskop microscope (Zeiss).

### Statistical analysis

Statistical analyses were performed using GraphPad Prism software6.0 (San Diego, CA, USA). Data were analyzed by using an unpaired Student’s t-test when comparing two experimental groups. And one-way ANOVA with Bonferroni’s post-hoc test was used when comparing more than two experimental groups. Statistics were calculated using two-way ANOVA Tukey’s post-hoc test when comparing more than two groups with two factors.

## RESULTS

### Apoptotic cells are preferentially localised to CD103+ve DCs coursing from the murine gut

In the mammalian gut wall there is large-scale spontaneous death by apoptosis of intestinal epithelial cells (IECs) which are marked by positive staining for non-specific esterase (NSE). However, in the murine system it was not yet known whether CD103+ve DCs played a significant role in removing apoptotic cells arising spontaneously, as inferred from seminal studies [7] of an analogous DC population in rats, known to transport apoptotic IECs to draining mesenteric lymph nodes (MLNs).

We prepared MLNs from healthy C57BL/6 mice housed in pathogen-limited conditions, selecting CD11c+ve DCs which were sorted for CD103 expression and stained by TUNEL for prior ingestion of cells containing apoptotic, cleaved double-stranded DNA. Inclusions of apoptotic material were exhibited 24.7-fold more frequently by CD103+ve DCs, when compared with CD103-ve DCs (Figure 1A). Because rapid degradation of DNA occurs in phagolysosomes, we also stained for the more stable IEC marker NSE, observing that NSE-positive inclusions were 8-fold more frequent in CD103+ve DCs (Figure 1B). Thus, when compared with CD103-ve MLN DCs, there was evidence that a much greater proportion of CD103+ve MLN DCs had spontaneously ingested apoptotic cells, including IECs, prior to reaching the draining lymph node.

**Figure 1.**
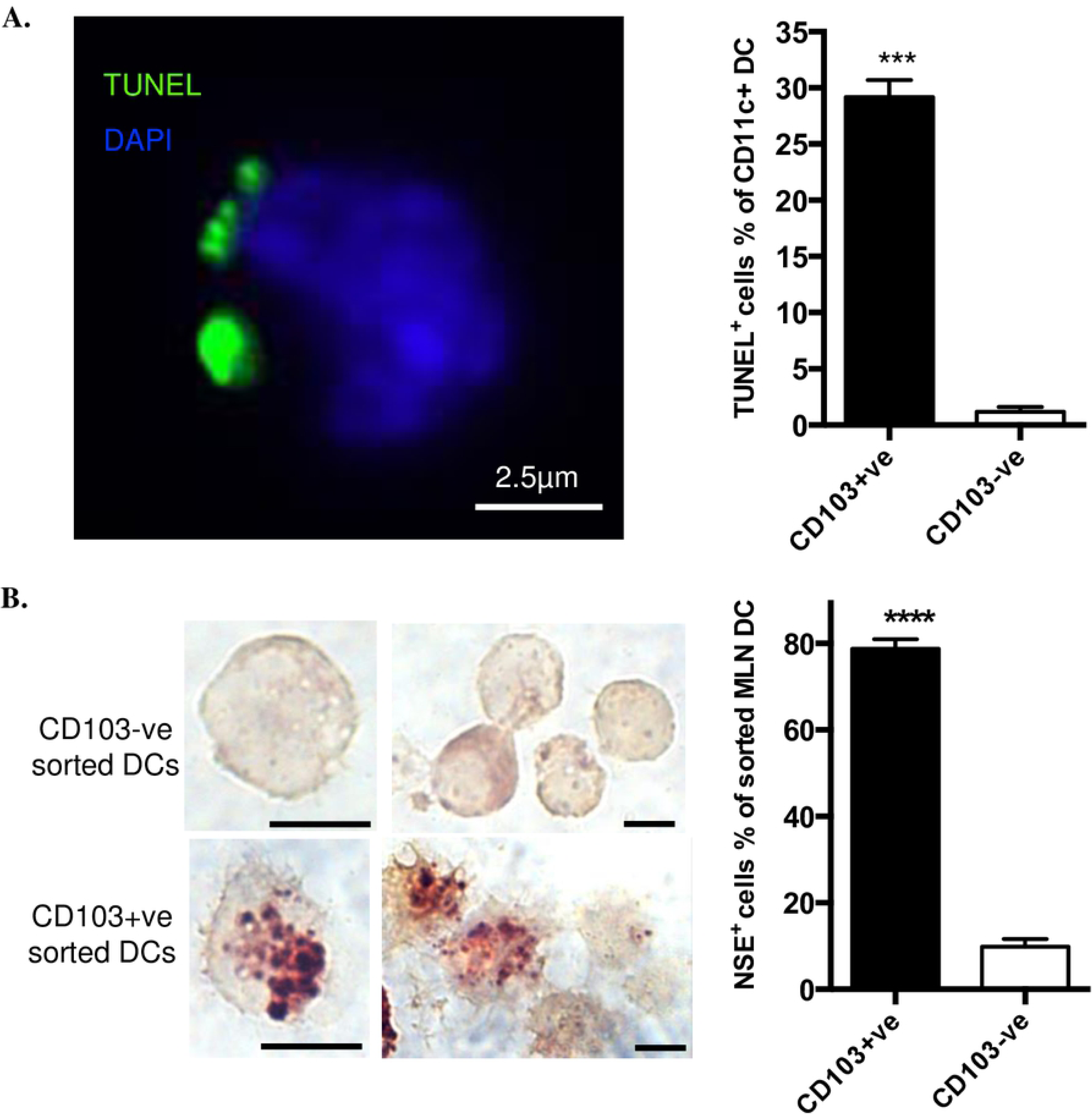
CD103+ve DCs from MLN contain apoptotic cell remnants. **A.** FACS-sorted CD103+ve DCs and CD103-ve DCs were prepared by affinity beads from pooled MLN from three mice and stained with TUNEL (green) and DAPI (blue). One typical image (left) shows TUNEL positive material associated with CD103+ve MLN DC. Chart (right) shows percentage of DCs bearing TUNEL positive material. **B.** Staining for the marker of intestinal endothelial cells, Non-Specific Esterase (NSE) was performed on CD11C+ve DCs isolated from MLNs. Scale bar, 5µm. All data were from four experiments with at least 200 cells counted for each experiment. Data show mean ± SEM of four individual experiments. *** p<0.001; **** p<0.0001; unpaired t test.

### Phagocytosis of apoptotic cells *ex vivo* by MLN DCs does not depend on CD103 expression

Preferential localisation *in vivo* of apoptotic cells to CD103+ve DCs isolated from MLNs (Figure 1) could reflect different capacity for phagocytosis of apoptotic cells by mature DC populations defined by CD103 expression. However, when we compared the capacity of isolated CD103+ve and CD103-ve MLN DCs to ingest apoptotic thymocytes *ex vivo*, no significant difference was observed (Figure 2A, 2B). This indicated that rather than greater phagocytosis of apoptotic IECs *in situ* being a passive consequence of mature MLN DC phagocytic capacity co-incidentally associated with the CD103+ve phenotype, phagocytosis of apoptotic cells during maturation of DC precursors might preferentially drive CD103 expression.

**Figure 2.**
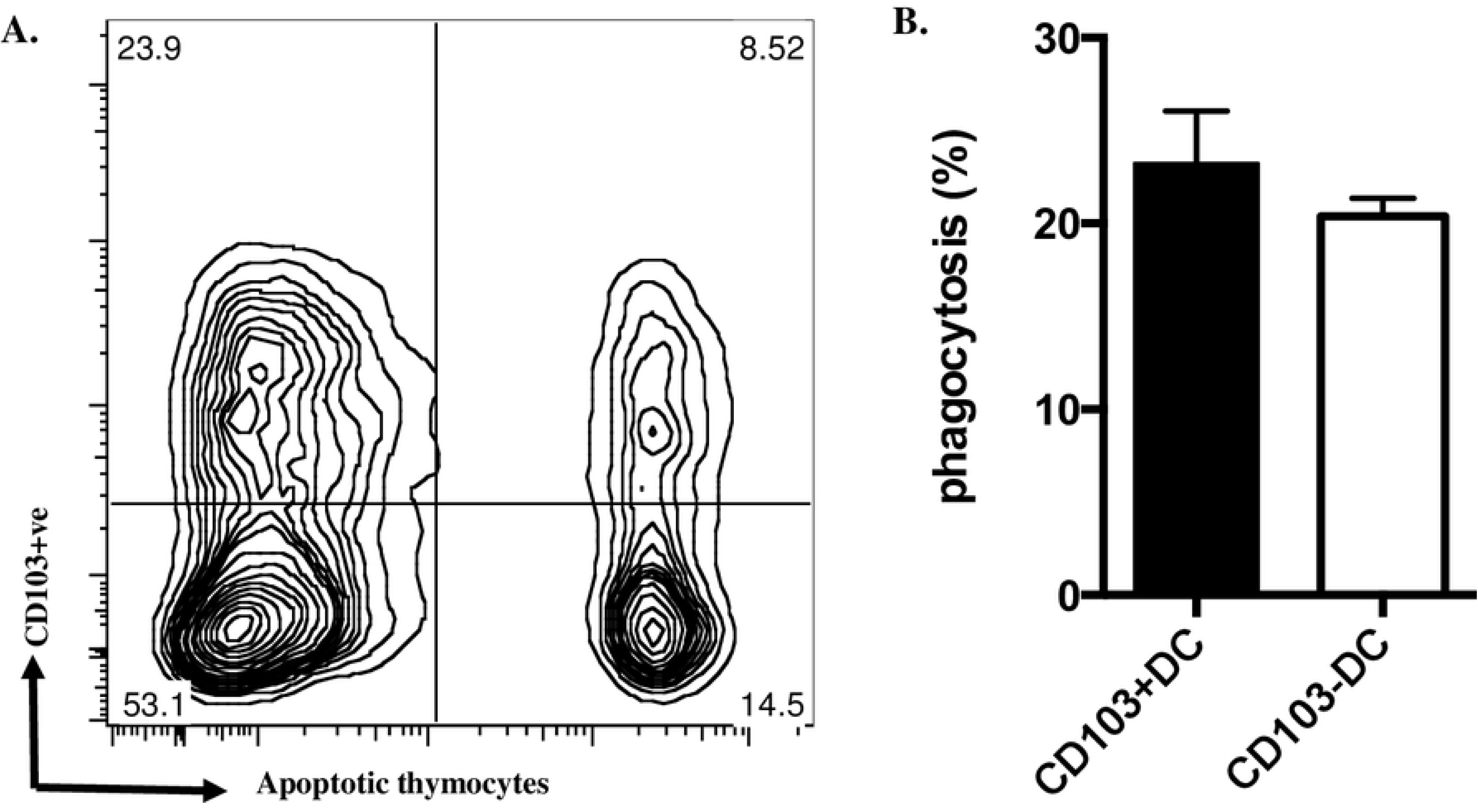
Phagocytosis of apoptotic cells *ex vivo* by MLN DCs does not depend on CD103 expression. **A.** MLN DCs co-cultured with fluorescence-labelled apoptotic thymocytes for 2h were stained for CD103 and assessed by flow cytometry; a typical plot is shown; note the similar proportion of DCs ingesting apoptotic cells in both CD103+ve and CD103-ve populations. **B.** from three such experiments. Data show mean ± SEM, no significant difference; unpaired t test.

### Apoptotic cells drive CD103 expression in myeloid DC precursors *in vitro*

To test possible effects of apoptotic cells on myeloid DC differentiation we created a model system *in vitro*, based on myeloid DC precursors isolated by a protocol closely similar to that described by del Rio *et al* and characterised as having the expected lineage-ve CD11c-ve CD103-ve phenotype [17]. In keeping with monocytic precursors of macrophages [18], such myeloid DC precursors bound but did not ingest either syngeneic apoptotic thymocytes or human apoptotic neutrophils (Figure 3A); the latter represent a particularly pure apoptotic cell “meal” with low (<2%) levels of secondary necrosis [14, 15]. When myeloid DC precursors were co-cultured with apoptotic human neutrophils at a 1:1 ratio, the relative proportion of DCs expressing CD103 at 7 days was increased by 53.2±9.04 % (mean±SEM, n=4) (Figure 3B). When co-cultured at varying DC:Ac (apoptotic cell) ratios, increasing numbers of apoptotic cells did not recruit more DC precursors into the CD103+ve DC population at 7 days (Figure 3C) but did increase the intensity of CD103 staining (Figure 3D). When apoptotic cells were added at different times after myeloid DC precursor culture was established (Figure 3E), recruitment of DCs into the CD103+ve population at 7d required co-culture with apoptotic cells prior to day 3, with maximal effect being observed when apoptotic cells were included from day 0, indicating that DC precursors rather than more mature DCs exhibited upregulation of CD103 when exposed to apoptotic cells. We confirmed (Figure 3F) that apoptotic cells induced DC precursor CD103 expression by qRTPCR analysis of mRNA for CD103 and the associated immunoregulatory marker, the β_8_ integrin chain, which is critical for myeloid DCs to activate TGF-β1 and elicit responses such as induction of Tregs [2, 3, 11, 19].

**Figure 3.**
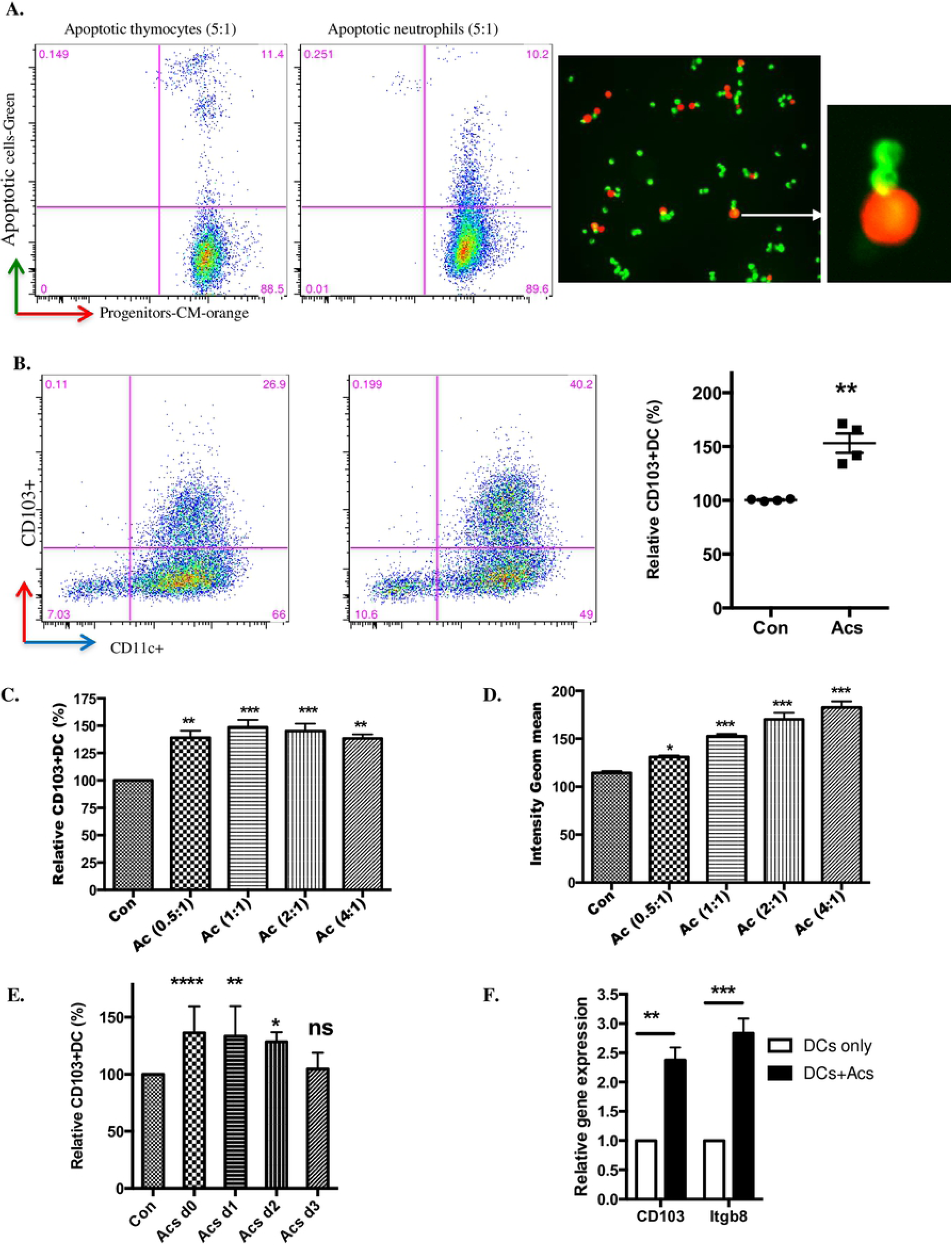
Coculture with apoptotic cells promotes CD103 expression by DC precursors. **A.** Freshly isolated DC precursors stained with CM-orange were co-cultured for 2hrs with apoptotic mouse thymocytes or apoptotic human neutrophils, both labelled with Cell Tracker Green. DC precursors binding apoptotic cells was measured by flow cytometry. Plots (left) show the gated CM-orange DC precursors with green apoptotic cells. Images (right) show DC precursors binding apoptotic thymocytes; note merged, yellow staining at the binding points. **B.** Selected bone marrow DC precursors cultured with apoptotic human neutrophils at the of 1:1 (Ac:DC) at day0 in the presence of GM-CSF. Cells were stained with CD11c-APC and CD103-PE after 7days. Flow Cytometry was performed and one plot representative of four is shown (left). The chart (right) shows relative CD103+ve DC percentage from four different experiments. **C, D.** DC precursors co-cultured with apoptotic cells in different ratios as indicated (Ac:DC) at day0. Relative CD103+ve DC percentage and geometric mean intensity were measured by flow cytometry after 7days. Data shows one experiment representative of three. **E.** DC precursors cultured with apoptotic cells at the ratio of 1:1 (Ac:DC) and apoptotic cells added at days 0, 1, 2 and 3. Relative CD103+ve DC (%) were analysed at day7. **F.** DC precursors cultured with apoptotic cells at the ratio of 1:1 (Ac:DC) at day0 and relative gene expression was measured by qPCR after 7 days. Data were from three different experiments. All data show mean ± SEM. * p<0.05; **p<0.01; *** p<0.001; **** p<0.0001; one-way ANOVA with Dunnet’s post-hoc test (B, C, D, E) or two-way ANOVA with Tukey’s post-hoc test (F).

### DC precursors co-cultured with apoptotic cells support enhanced induction of Tregs

Next, we sought a functional effect of apoptotic cell-directed induction of CD103 expression in DC precursors differentiating *in vitro*. DC precursors cultured for 7 days with (or without, as a control) apoptotic cells were then co-cultured for a further 4 days with murine CD4+ve T cells bearing a FoxP3/GFP transgene marker as described [3, 20]. When compared with DCs that had not been exposed to apoptotic cells, induction of FoxP3+ve T cells by either latent TGF-β1 at 2ng/ml or latent TGF-β1 at the same concentration plus retinoic acid at 100nm was significantly enhanced by DC precursor cultures that had been exposed to apoptotic cells from day 0 (Fig 4A). Furthermore in standard assays of antigen-driven T cell proliferation, suppression of T cell activation was greater with T cell populations induced by DCs exposed to apoptotic cells when compared with DCs that had not (Fig 4B). Thus, apoptotic cell direction of enhanced CD103 expression was indeed correlated with enhanced immunoregulatory function of DCs generated *in vitro*.

**Figure 4.**
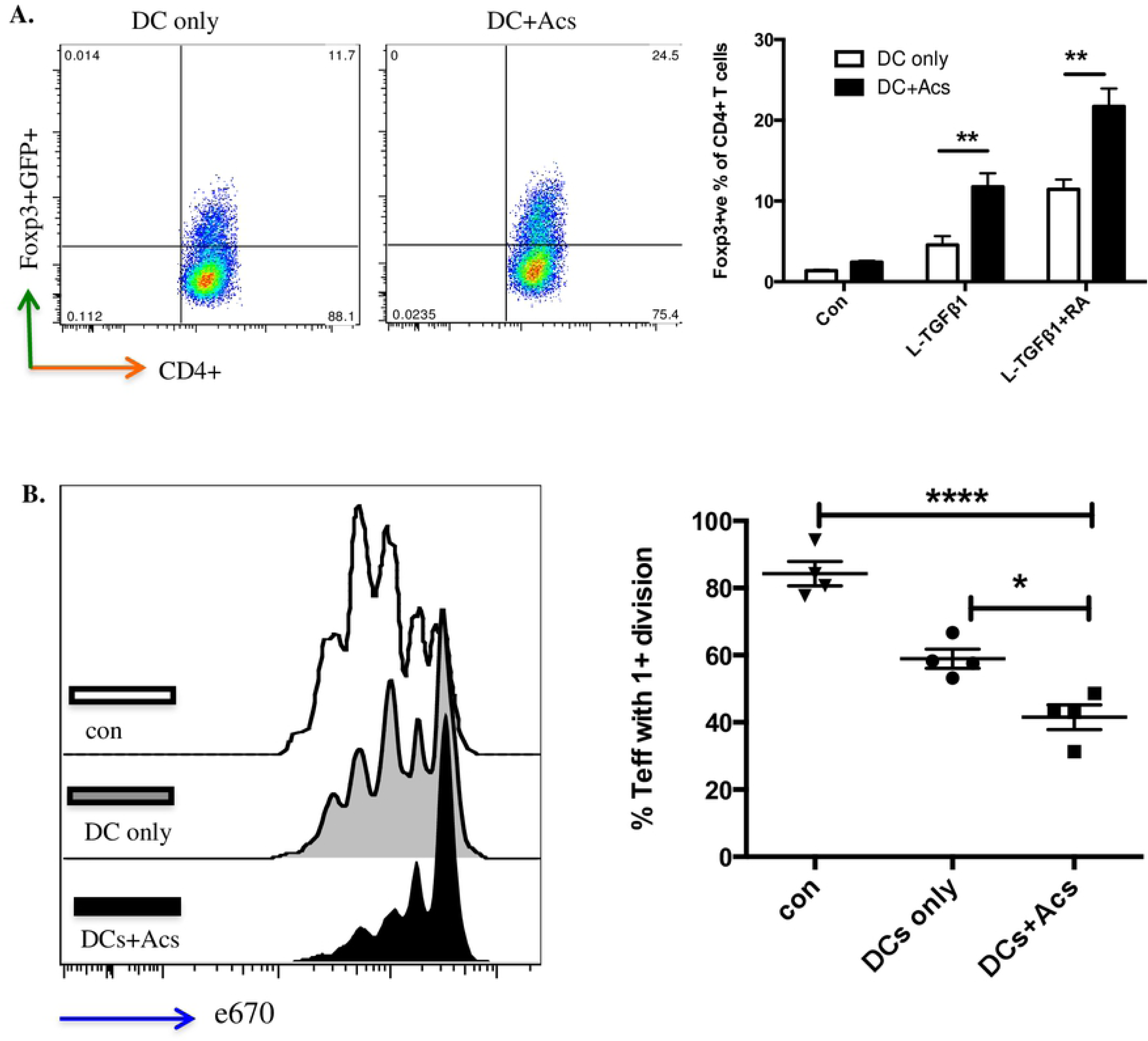
CD103+ve DCs induced by apoptotic cells enhance functional Treg development. **A.** BMDCs differentiated from DC precursors co-cultured with naïve CD4+ve GFP-ve FoxP3-ve T cells in the presence of 2ng/ml latent TGF-β1 ± 100nm retinoic acid (RA). The percentage of FoxP3+ve T cells was determined at day 5 by flow cytometry (left); chart (right) collates data from three experiments. **B.** Naïve CD4+ve CD25-ve effector T cells from OTII mouse spleen were labelled with eFluor® 670 (e670) and co-cultured at 1:1 ratio with CD4+ T cells only (con); CD4+ T cells co-cultured with DCs in the presence of 2ng/ml latent TGF-β1 + 100nm RA for 5 days (DCs only); CD4+ T cells co-cultured with DCs exposed to apoptotic cells in the presence of 2ng/ml latent TGF-β1 + 100nm RA (DC+Acs) for 5 days. pOVA pulsed DCs were added in all wells to stimulate effector T cells. Effector T cell proliferation was determined after 72hrs by flow cytometry; plots show one experiment representative of three. Data show mean ± SEM of four replicate wells. **p<0.01; *** p<0.001; two-way ANOVA with Tukey’s post-hoc test (A); one-way ANOVA with Dunnet’s post-hoc test (B).

### DC precursor α_v_ integrin orchestrates binding of apoptotic cells with activation of TGF-β1

Previously we have shown by bone marrow transplantation and other strategies that in α_v_-tie2 mice [9], in which a tie2-CRE construct deletes floxed α_v_, the effects on DC function are due to deletion of α_v_ in the myeloid line. Having confirmed that myeloid DC precursors from α_v_-tie2 mice lacked α_v_ expression (Fig 5A), we then undertook an assay of the binding of apoptotic cells by myeloid DC precursors (3hrs), observing that this was reduced by about half (Fig 5B), in keeping with many studies of the complex multi-molecular mechanisms underlying myeloid phagocyte recognition of apoptotic cells [8, 16, 21–24], in which phagocytosis of apoptotic cells was reduced by about half where a putative phagocyte receptor for apoptotic cells, such as α_v_ was deleted [9]. This indicated that DC precursor α_v_ integrin is necessary for maximally efficient binding of apoptotic cells and that other mechanisms, not examined further in this study, contributed to binding. However, α_v_ deficiency completely abrogated the capacity of apoptotic cells and/or latent TGF-β1 (administered at day 0) to increase myeloid DC precursor CD103 expression at day 7 (Fig 5C). The dependence of such increased expression upon active TGF-β1 was confirmed by the capacity of blocking antibody to inhibit direction of increased DC precursor CD103 expression by apoptotic cells ± latent TGF-β1 irrespective of whether α_v_ was expressed (Fig 5C). These observations led us to seek direct evidence that α_v_ integrins and latent TGF-β1 might be brought together at points where DC precursors contact apoptotic cells.

**Figure 5.**
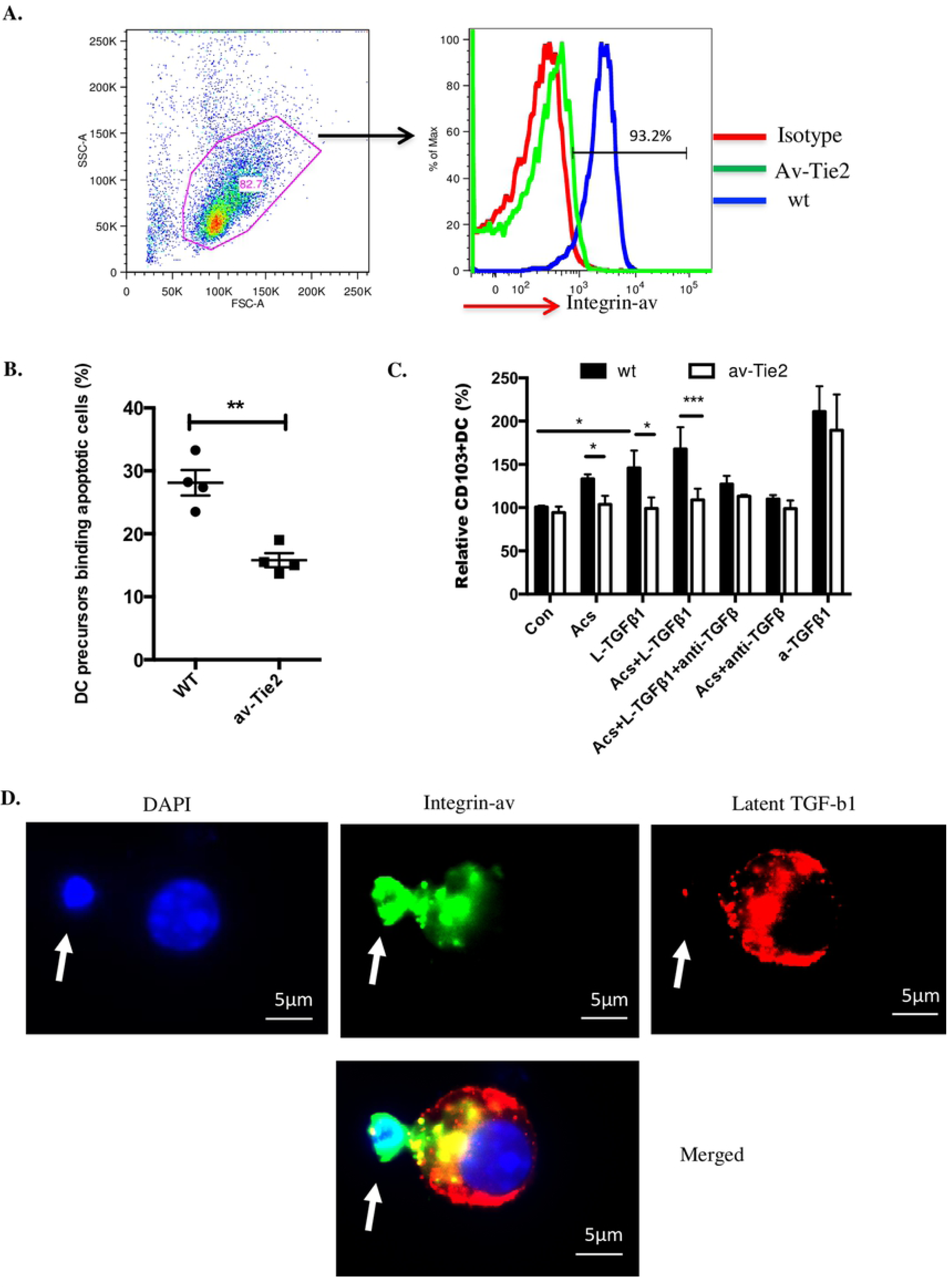
DC precursor v integrin orchestrates binding of apoptotic cells, activation of TGF-β1 and enhanced CD103 expression. **A.** DC precursors from wt and α_v_-tie2 mice were stained with v-integrin-PE antibody and flow cytometry performed; plot representative of three experiments. **B.** DC precursors (stained with CM-orange) from wild type or α_v_-Tie2 mice cultured with mouse apoptotic thymocytes (stained with Cell Tracker Green) at a ratio of 5:1 (Ac:DC). Binding was measured by flow cytometry after three hours. Data were accumulated from two experiments with four mice. **C.** CD103 expression by DCs derived from DC precursors cultured in serum free X-vivo medium in presence of GM-CSF. Conditions were DC precursor only (Con); DC precursor cultured with apoptotic cells at 1:2 ratio (Acs); DC precursor cultured with 2ng/ml latent TGF-β1 (L-TGFβ1); DC precursor cultured with apoptotic cells and 2ng/ml latent TGF-β1 (Acs+L-TGFβ1); DC precursor cultured with apoptotic cells and 2ng/ml latent TGF-β1 and 2μg/ml anti-TGF-β (Acs+L-TGFβ1+anti-TGFβ); DC precursor cultured with apoptotic cells and 2μg/ml anti-TGF-β (Acs+anti-TGFβ); DC precursor cultured with 0.2ng/ml active-TGF-β1 (a-TGFβ1). Relative percentage of CD103+ve DCs were measured by flow cytometry after 7days. Data were from three different experiments with three mice per group. **D.** DC precursors from wild type mice cultured with apoptotic thymocytes at a ratio of 1:5 for three hours. Cells were stained with integrin α_v_ (green), LAP (latent TGF-β1; red) and DAPI. White arrow points an apoptotic cell binding to a DC precursor. All data show mean ± SEM. * p<0.05; **p<0.01; *** p<0.001; **** p<0.0001; unpaired t test (B) or two-way ANOVA with Tukey’s post-hoc test (C).

We employed three colour fluorescence microscopy to examine non-permeabilised DC precursors at 4oC after co-culture with apoptotic thymocytes at 37 oC for 3 hours, consistently finding that 31.23 ± 3.706 (%) (mean ± SEM, n= 3) precursors bound apoptotic cells. Staining for α_v_ integrin was often intense at points of contact, with α_v_ -positive DC processes partially surrounding bound apoptotic cells in cup-like structures (Figure 5D); α_v_ staining was present at 91.8 ± 5.7 (%) (mean ± SEM, n= 3) of points of contact. Low intensity staining for latent TGF-β1 was associated with the surface of both DC precursors and apoptotic thymocytes, but high intensity staining co-localised with α_v_ –positive points of contact in 72.2 ± 1.5 (%) (mean ± SEM, n=3). For obvious reasons we could not seek co-localisation of α_v_ staining with latent TGF-β1 staining with DC precursors from α_v_–tie2 mice, but in the small proportion of α_v_-negative points of contact between wild type DC precursors and apoptotic cells we never observed staining for latent TGF-β1.

## DISCUSSION

Myeloid phagocyte populations present in the healthy mammalian gastrointestinal tract are complex and the subject of ongoing investigation in many laboratories. Nevertheless, CD103+ve myeloid DCs have been characterised in detail in the mouse, with compelling evidence for key roles in keeping the healthy gut free of inflammation, by induction of Tregs and other mechanisms dependent on TGF-β1 activation [2, 3, 5, 19, 25]. In this report, we demonstrate that CD103+ve DCs isolated from the mesenteric lymph nodes (MLNs) of healthy mice exhibited extensive evidence of prior ingestion of IECs undergoing spontaneous apoptosis whereas, depending on the straining technique used to identify previously ingested apoptotic IECs, CD103-ve MLN DCs were 8-24 fold less likely to do so. Because the *ex vivo* capacity to ingest apoptotic cells was similar for both CD103+ve and CD103-ve MLN DCs, we sought evidence that phagocytosis of apoptotic cells might drive CD103-ve DC precursors towards the immunoregulatory phenotype of mature CD103+ve DCs. In an *in vitro* model system apoptotic cells did indeed direct enhanced expression by co-cultured lineage-ve CD103-ve myeloid DC precursors of the immunoregulatory CD103+ve β_8_+ve DC phenotype. These immunoregulatory CD103+ve DC also enhanced induction of functional Tregs *in vitro*. However, when myeloid DC precursors were isolated from mice deleted for α_v_ integrin in the myeloid line, DC precursors exhibited reduced binding of apoptotic cells and complete loss of enhancement of CD103 expression by apoptotic cells and/or latent TGF-β1; a role for α_v_ integrin-mediated activation of latent TGF-β1 was confirmed by the capacity of active TGF-β1 to increase CD103 expression in DC precursors from α_v_-tie2 mice and wild type mice. Because we observed co-localisation of DC precursor α_v_ integrin and latent TGF-β1 at points of contact with bound apoptotic cells, we conclude that immature CD103-ve DC precursors can deploy α_v_ integrins to capture apoptotic cells and co-ordinately activate latent TGF-β1. This results in differentiation of CD103-ve DC precursors towards a mature, immunoregulatory CD103+ve DC phenotype. Thus, a hitherto unrecognised consequence of myeloid phagocyte binding and uptake of apoptotic cells may be to programme DCs to promote the tonic suppression of inflammation in the healthy gut.

Evidence of prior ingestion of apoptotic cells by dendritic cells coursing from the gut to mesenteric lymph nodes was first described in the rat by MacPherson’s group [7]. The OX41-ve DC subset in MLNs was found to have preferentially ingested apoptotic IECs and was speculated to have a role in inducing and maintaining self-tolerance [7]. However, despite functional similarities between OX41-ve MLN DCs in rats and CD103+ve MLN DCs in mice, the current study is the first to confirm in healthy unmanipulated mice that CD103+ve MLN DCs do indeed exhibit extensive evidence of prior ingestion of apoptotic cells, particularly IECs. Our data confirm the physiological relevance of elegant transgenic studies in which IECs induced to die by apoptosis were tracked as being ingested by CD103+ve DCs exhibiting anti-inflammatory and immunosuppressive gene expression signatures [1].

A key observation in the current studies of MLN DCs was that similar capacity to ingest apoptotic cells *ex vivo* was observed for CD103-ve and CD103+ve CD11c+ve DCs isolated from MLNs, whereas only the latter exhibited extensive evidence of prior ingestion of apoptotic cells *in vivo*. This raised the possibility that differentiation of immature myeloid DCs towards a CD103+ve β_8_+ve immunoregulatory phenotype might be directed by the effects of contact with apoptotic cells upon immature CD103-ve myeloid DCs. The *in vitro* data reported here strongly support the likelihood of contact with apoptotic cells being a major factor, along with TGF-β1 activation, in driving acquisition by immature CD103-ve DCs of the CD103+ve phenotype associated with the capacity to generate functionally active Tregs.

One way of testing the dependence of differentiation of gastrointestinal tract myeloid DCs upon binding and ingestion of apoptotic cells would be to characterise mechanisms by which immature DCs interact with apoptotic cells and elaborate active TGF-β1. In the current study we demonstrate, using mice deficient for α_v_ in the myeloid line, that *in vitro* myeloid DC precursor binding of apoptotic cells is partially dependent upon DC precursor α_v_ integrin. Furthermore, DC α_v_ integrin is essential for the capacity of apoptotic cells and/or latent TGF-β1 to enhance CD103 expression by DC precursor populations. These *in vitro* data would predict that mice deficient for α_v_ in the myeloid line would exhibit *in vivo* (a) reduced capacity of myeloid phagocytes to bind and ingest apoptotic cells; (b) diminished numbers of CD103+ve DCs in MLNs; (c) reduced numbers of Tregs in MLNs; and (d) spontaneous inflammation in the gut. All of these characteristics were exhibited by α_v_-tie2 mice in which the deleterious effects of α_v_ deficiency were shown to be due to effects on myeloid phagocytes [9].

Therefore, we propose that in the gut a hitherto unrecognised consequence of the interaction between immature DCs and apoptotic cells arising spontaneously is the beneficial “programming” of maturing DCs towards the CD103+ve β_8_+ve phenotype that is known to keep gut inflammation in check by tonic suppressive mechanisms. The capacity of apoptotic cells to suppress inflammation and enhance active TGF-β1 elaboration was initially studied in models of resolving acute inflammation [16, 26-–28] but the current study emphasises that apoptotic cells may pro-actively prevent inflammation in addition to suppressing active inflammation, in both cases by altering the “programming” of myeloid phagocytes.

Further work is obviously needed to dissect in detail the mechanisms by which binding of apoptotic cells by DC precursors directs subsequent DC differentiation. Nevertheless, the current study supports the hypothesis that contact with apoptotic cells allows clustered DC surface α_v_ integrin to orchestrate the formation of multimolecular complexes that can capture and activate latent TGF-β1. Such future work should address which of a number of potential “bridging” molecules allow DC surface α_v_ integrin to bind apoptotic cells. Of those previously implicated [16, 22, 23, 29], it is intriguing that molecules such as thrombospondin are capable of activating latent TGF-β1[30]. Furthermore, although enhancement of DC precursor CD103 expression by apoptotic cells and/or latent TGF-β1 was completely dependent upon DC expression of α_v_, the data suggest that other DC surface receptors are probably involved in apoptotic cell capture, because α_v_ deficiency did not completely eliminate the capacity of DC precursors to bind apoptotic cells; for example, candidate receptors such as *Mer* are known to co-operate with α_v_ [31]. Indeed, the signalling mechanisms elicited within DC precursors that deploy α_v_ integrin to bind apoptotic cells await characterisation. Lastly, because careful microscopy by Blander’s group showed that DCs already expressing CD103 can, in a gut stressed by induced large-scale apoptosis, select and bind apoptotic cells, future work should also seek to demonstrate interaction *in vivo* between DC precursors and apoptotic cells. For example, means should be sought to determine whether lineage-ve CD11c-ve CD103-ve DC precursors do closely appose to IECs dying by apoptosis in the gut wall, but such studies are beyond the scope of the current work.

To conclude, we report that murine CD103+ve MLN DCs, known to be important in constitutive suppression of gut inflammation, do indeed exhibit extensive evidence of prior ingestion of IECs dying spontaneously by apoptosis. Furthermore, we demonstrate *in vitro* that exposure to apoptotic cells directs the differentiation of CD103-ve myeloid DC precursors towards the mature CD103+ve β_8_+ve myeloid DC phenotype capable of inducing Tregs. The mechanisms responsible involve DC precursor α_v_ integrin, which orchestrates the binding of apoptotic cells with the activation of latent TGF-β1 and enhancement of the immunoregulatory CD103+ve phenotype. These data, taken with earlier observations of spontaneous gut inflammation in animals lacking α_v_ in the myeloid line, suggest that a newly identified consequence for myeloid phagocyte interaction with apoptotic cells is programming of DCs to prevent inflammation threatened by micro-environmental stimuli.

## Acknowledgment

We thank Stephen Anderton and Richard A. O’Connor for suppling FoxP3 reporter mice and CD45.1 OTII mice. We thank the QMRI Flow Cytometry and Cell Sorting Facility for their excellent technical assistance. Eleanor Bonikowski and Ruth MacInnes provided invaluable secretarial help with the manuscript. We also thank Professor Kev Dhaliwal and Professor Neil Henderson for supporting animal experiments.

## Author Contributions

Ailiang Zhang, Helena Paidassi, Adam Lacy-Hulbert, and John Savill conceived the study and edited the manuscript; Ailiang Zhang and Helena Paidassi performed the experiments; Ailiang Zhang and John Savill wrote and reviewed the manuscript.

